# Genomics of *Cryptococcus neoformans*

**DOI:** 10.1101/356816

**Authors:** PM Ashton, LT Thanh, PH Trieu, D Van Anh, NM Trinh, J Beardsley, F Kibengo, W Chierakul, DAB Dance, LQ Hung, NVV Chau, NLN Tung, AK Chan, GE Thwaites, DG Lalloo, C Anscombe, LTH Nhat, J Perfect, G Dougan, S Baker, S Harris, JN Day

## Abstract

*C. neoformans* var. *grubii* (*C. neoformans*) is an environmentally acquired pathogen causing 181 000 HIV-associated deaths each year. We used whole genome sequencing (WGS) to characterise 699 isolates, primarily *C. neoformans* from HIV-infected patients, from 5 countries in Asia and Africa. We found that 91% of our clinical isolates belonged to one of three highly clonal sub-clades of VNIa, which we have termed VNIa-4, VNIa-5 and VNIa-93. Parsimony analysis revealed frequent, long distance transmissions of *C. neoformans*; international transmissions took place on 13% of VNIa-4 branches, and intercontinental transmissions on 7% of VNIa-93 branches. The median length of within sub-clade internal branches was 3-6 SNPs, while terminal branches were 44.5-77.5 SNPs. The short median internal branches were partly driven by the large number (12-15% of internal branches) of polytomies in the within-sub-clade trees. To simultaneously explain our observation of no apparent molecular clock, short internal branches and frequent polytomies we hypothesise that *C. neoformans* VNIa spends much of its time in the environment in a quiescent state, while, when it is sampled, it has almost always undergone an extended period of growth. Infections with VNIa-93 were associated with a significantly reduced risk of death by 10 weeks compared with infections with VNIa-4 (Hazard Ratio = 0.45, p = 0.003). We detected a recombination in the mitochondrial sequence of VNIa-5, suggesting that mitochondria could be involved in the propensity of this sub-clade to infect HIV-uninfected patients. These data highlight the insight into the biology and epidemiology of pathogenic fungi which can be gained from WGS data.

## Intro

*Cryptococcus neoformans* is an opportunistic fungal pathogen which primarily affects people with cell mediated immune defects, particularly those living with HIV. There are an estimated 223 100 incident cases of cryptococcal meningitis per year in HIV patients with CD4 counts of less than 100 cells per µl, resulting in 181 100 deaths (Rajasingham et al. 2017). *C. neoformans* var. *grubii* (hereafter *C. neoformans*), one of two varieties of *C. neoformans*, accounts for the vast majority of cryptococcal meningitis cases globally, and particularly in the tropical and sub-tropical regions which bear the heaviest disease burden (Rajasingham et al. 2017; Park et al. 2009).

The population structure of *C. neoformans* consists of at least three lineages, VNI, VNII and VNB. Two of these, the frequently isolated VNI and the rarely observed VNII, are clonal and globally distributed (Litvintseva et al. 2006; Khayhan et al. 2013; Ferreira-Paim et al. 2017) while VNB is very diverse but rarely isolated outside sub-Saharan Africa (Litvintseva et al. 2006) and South America (Andrade-Silva et al. 2018). Sequencing of strains from patients with relapsed disease has indicated that microevolution occurs during infection, with typically 0-6 SNPs occurring over a median relapse period of 146 days (Chen et al. 2017). Other studies have described a broad view of the three main molecular types, VNI, VNII and VNB, analysing 150-400 total isolates, and placing clinical isolates into the context of environmental strains (Desjardins et al. 2017; Rhodes et al. 2017; Vanhove et al. 2017). Within VNI, three distinct, but still recombining, sub-lineages have been identified, two of which (VNIa and VNIb) are globally distributed, while VNIc is limited to southern Africa. Genomic data has revealed that VNI and VNII to have more recent migrations than VNB, with nearly clonal isolates found in disparate geographic regions (Rhodes et al. 2017), although this has not yet been investigated on a fine scale.

So far, our understanding of the population structure of *C. neoformans* in the Asia & Pacific region, the second highest prevalence region after sub-Saharan Africa (Rajasingham et al. 2017), has been based upon low resolution methods such as MLST and AFLP (Day et al. 2011; Thanh et al. 2017; Simwami et al. 2011; Khayhan et al. 2013; Kaocharoen et al. 2013; Hiremath et al. 2008; Day et al. 2017). These data show that *C. neoformans* in Southeast Asia is highly clonal, with considerable gene flow between countries within the region, and less connectivity with other continents (Khayhan et al. 2013). Recently, the first study focussing on whole genome data from the region has been reported, which identified 165 Kbp of sequence specific to ST5 (Day et al. 2017), a sequence type seen more frequently n HIV uninfected patients, the majority of whom have no identified underlying immune-suppression (Day et al. 2011, 2017). The predilection of ST5 to infect HIV uninfected patients is not the only reported association between a *C. neoformans* lineage and a clinical phenotype. Infections with VNB (Beale et al. 2015) and VNI ST93 (Wiesner et al. 2012) have been reported to have worse outcomes in HIV infected patients in southern Africa and eastern Africa, respectively.

Production of *C. neoformans* spores is thought to be vital to the organism’s virulence, as the spores, alongside desiccated yeast cells are the likely infectious propagule (Velagapudi et al. 2009). There are two known mechanisms which can result in the generation of *C. neoformans* spores –heterothallic mating and homothallic fruiting. Both processes involve meiosis resulting in recombination and other large scale genomic changes such as aneuploidy (Lin and Heitman 2006; Ni et al. 2013; Lin et al. 2005). While our direct understanding of spore production in *C. neoformans* comes entirely from the laboratory, evidence of the processes occurring naturally have mostly come indirectly from population genetics (Litvintseva et al. 2006; Hiremath et al. 2008). Previously, we have undertaken several prospective, descriptive and randomised controlled intervention trials in Southeast Asia and East/Southeast Africa. Here, we used whole genome sequence analysis of 699 *Cryptococcus* isolates to describe the population structure of *C. neoformans* causing disease in these populations, in high resolution, and combine this information with metadata from these trials to relate this to disease phenotype.

## Results

We sequenced 699 *Cryptococcus* species complex isolates from Vietnam (n = 441), Laos (n = 73), Thailand (n = 40), Uganda (n = 132) and Malawi (n = 13). Of these, 682 were *C. neoformans*, 12 were *C. gattii* and 5 (all from Uganda) were putative hybrids between *C. neoformans* and *C. deneoformans*. There were 696 clinical isolates from 695 patients, and 3 environmental isolates from Vietnam. All environmental isolates were *C. neoformans*. There were 618 isolates from HIV infected patients and 78 from HIV uninfected patients. Of the 682 *C. neoformans* there were 681 isolates with mating type alpha and 1 isolate from Vietnam with mating type a.

### Whole genome sequencing of VNI

Six hundred and seventy eight (99.4%) of our *C. neoformans* isolates were VNI; four were VNII (Supplementary Figure 1, Supplementary Table 1). To provide context for our isolates, all 185 VNI genomes sequenced by Desjardins *et al*. (160 clinical, 25 environmental, full details available in Supplementary Table 1) were included in subsequent phylogenetic analyses. We ensured technical comparability of our methods of phylogenetic analysis with those of Desjardins *et al.* by comparing our results for the Desjardins data with their reported results (Supplementary Figure 2).

A phylogenetic tree (Figure 1) was derived from the 325812 variant positions in the core genome of the 863 *C. neoformans* VNI. Of the novel *C. neoformans* isolates presented here, 668 were VNIa (98.5%), 10 were VNIb (1.5%); none were VNIc. Figure 1 shows that the population structure of VNIa is dominated by three common and highly clonal sub-clades, while VNIb and VNIc are more heterogenous. VNIa, VNIb and VNIc isolates were isolated from 14, 10 and 2 countries on 5, 6 and 1 continent(s), respectively (Supplementary Tables 2 & 3). VNIa was predominant, accounting for 548 of 549 (99.8%) isolates in Asia and 163 of 274 (59.5%) strains in Africa. When isolates from Botswana, an established outlier in terms of *Cryptococcus neoformans* diversity, were excluded, the proportion of VNIa isolates in Africa was 84.3% (134 out of 159) of all VNI isolates. The H99 reference genome belonged to VNIb.

**Figure 1:**
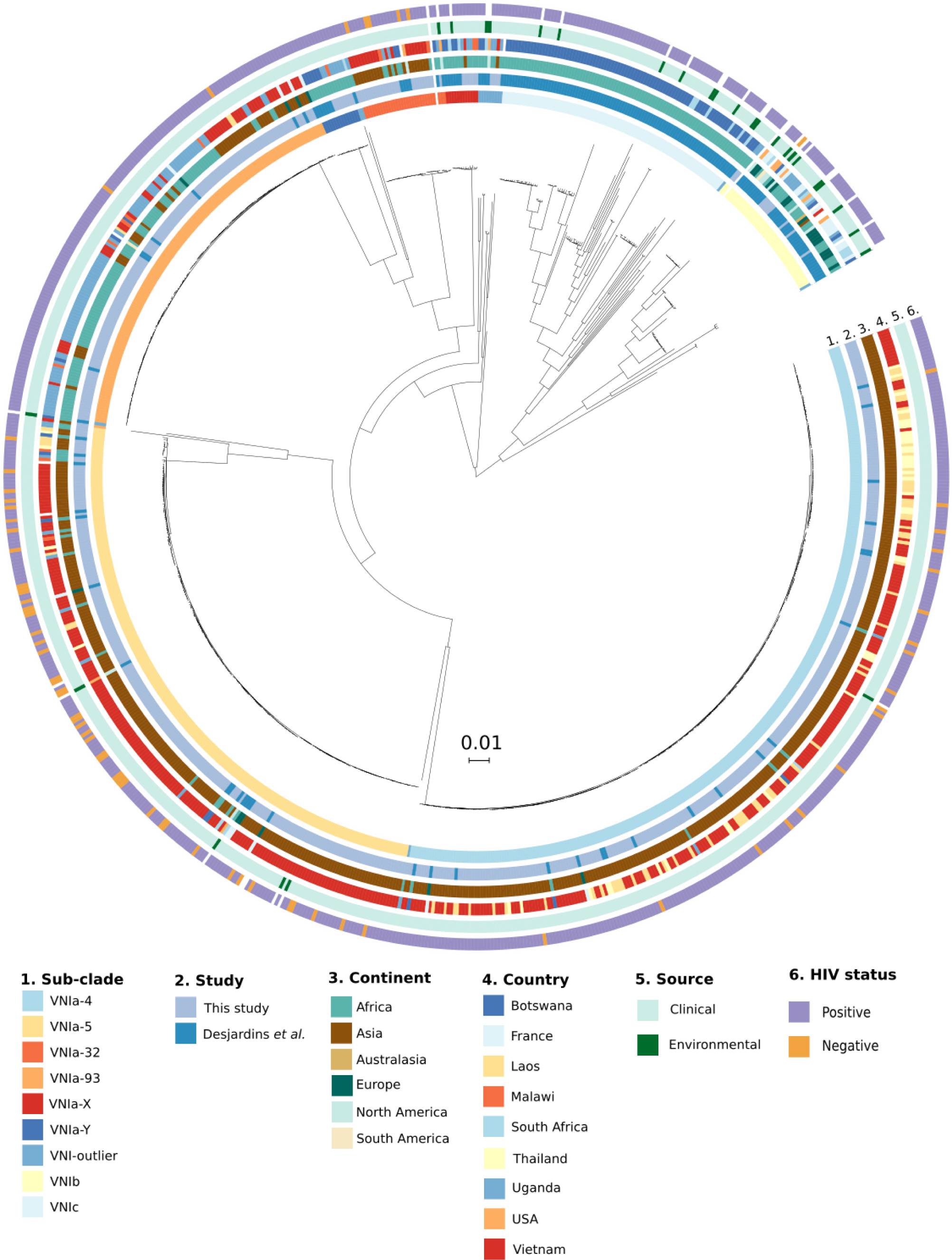
A whole genome SNP phylogeny of all VNI in this study and Desjardins et al

Nine distinct clusters were identified using PCA and K-means clustering (Supplementary Figure 3). We extended the naming scheme of Desjardins *et al*. to refer to the sub-clades within VNIa as VNIa-4, VNIa-5, VNIa-93 and VNIa-32 after the predominant MLST sequence type in each clade. Two clusters contained only isolates with novel STs, which we refer to as VNIa-X and VNIa-Y. The previously described VNIb and VNIc lineages were also identified as distinct clusters. The remaining polyphyletic VNI isolates which did not fall into any PCA cluster we grouped together into VNI-outlier. The number of each lineage isolated from HIV positive patients from each country are presented in Table 1.

**Table 1:** The frequency of isolation of each VNIa sub-clade from HIV positive patients in each country from both this study and Desjardins et al

| Country | VNIa-4 | VNIa-5 | VNIa-93 | VNIc | VNIb | VNIa-32 | VNIa-Y | VNIa-X | VNIa-outlier | Total |
| --- | --- | --- | --- | --- | --- | --- | --- | --- | --- | --- |
| Vietnam | 175 | 129 | 44 |  | 1 | 15 |  | 1 | 1 | 366 |
| Uganda | 2 | 8 | 84 |  | 10 | 3 | 7 | 3 | 5 | 122 |
| Botswana | 3 | 5 | 3 | 74 | 2 | 3 | 6 | 3 | 3 | 102 |
| Laos | 57 | 6 | 2 |  |  |  |  |  |  | 65 |
| Thailand | 38 | 4 |  |  |  |  |  |  |  | 42 |
| France | 2 | 4 | 4 |  | 15 |  |  |  |  | 25 |
| S. Africa | 1 | 1 |  | 6 | 6 |  |  | 2 | 1 | 17 |
| Malawi |  | 3 | 5 |  |  | 2 | 1 | 2 |  | 13 |
| Togo |  |  |  |  | 2 |  |  |  |  | 2 |
| India |  |  |  |  |  | 1 |  |  |  | 1 |
| Brazil |  |  | 1 |  |  |  |  |  |  | 1 |
| Argentina |  |  |  |  | 1 |  |  |  |  | 1 |
| Australia |  |  |  |  | 1 |  |  |  |  | 1 |
| USA |  | 1 |  |  |  |  |  |  |  | 1 |
| China |  | 1 |  |  |  |  |  |  |  | 1 |
| Japan |  | 1 |  |  |  |  |  |  |  | 1 |
| Tanzania |  |  |  |  |  |  |  |  | 1 | 1 |
| **Total** | **278** | **163** | **143** | **80** | **38** | **24** | **14** | **11** | **11** | **762** |

While each country had a dominant or, in the case of Vietnam, co-dominant sub-clade(s), there were minority sub-clades present in every country analysed (Supplementary Figure 4). For example, VNIa-93, the dominant lineage in Uganda, was also present in Vietnam (12%). Similarly, Uganda and Botswana had low prevalence of typically Southeast Asian sub-clades such as VNIa-4 (Uganda = 1.6%, Botswana = 2.9%) and VNIa-5 (Uganda = 6.5%, Botswana = 4.9%).

### Phylogenetic analysis of sub-clades within VNIa

We performed fine-scale genomic epidemiological analyses of VNIa for every sub-clade with at least 50 isolates from this study, i.e. VNIa-4, VNIa-5 and VNIa-93. These sub-clades accounted for 89.3% of the total isolates in our study, with VNIa-4 accounting for 41%, VNIa-5 for 29% and VNIa-93 for 20%. To maximise the phylogenetic resolution within these sub-clades, within sub-clade reference genomes were generated using PacBio sequencing (available via FigShare doi:10.6084/m9.figshare.6060686). The median SNP distance of the VNIa-4, VNIa-5 and VNIa-93 strains to the within sub-clade reference genome was 277 (Standard Deviation (SD) = 142), 338 (SD = 236) and 361 (SD = 44) SNPs, compared with 47619 (SD = 196), 46218 (SD = 245) and 48763 (SD = 262) to the H99 reference genome.

### Recombination Within Sub-Clades

Before deriving per sub-clade phylogenies from which genomic-epidemiological characteristics can be inferred, we quantified the extent to which recombination plays a role in generation of diversity within sub-clades. Recombination within sub-clades was investigated by assessing the degree of linkage disequilibrium (LD). LD was assessed for all within sub-clade SNPs with a minor allele frequency of 0.1 or greater. There was limited decay of LD as assessed by R2 generated by vcftools (Danecek et al. 2011), indicating minimal ongoing recombination (Supplementary Figure 5).

### Isolates from disparate geographical locations are interspersed within the sub-clade phylogenies

One of the most striking patterns observed in the per-sub-clade phylogenies is the interspersion of isolates from different countries and different continents throughout the phylogeny (see Figures 2 (A), (B) and (C)), indicating frequent international and intercontinental transmissions. We used parsimony analysis to quantify the minimum number of international transmission events which explain the current geographic distribution of strains. VNIa-4 had the largest number of international transmission events as a proportion of total internal branches (95% CI in parentheses, VNIa-4 = 13% (11-16%), VNIa-5 = 8% (6-11%), VNIa-93 = 10% (7-14%)), while VNIa-93 had the highest proportion of intercontinental branches (VNIa-4 = 1% (0-2%), VNIa-5 = 5% (3-7%), VNIa-93 = 7% (5-10%)).

### Notable within sub-clade phylogenetic features

A striking feature of the within sub-clade phylogenies is the combination of long terminal branch lengths and short internal branches. The median number of SNPs represented by the internal branch lengths compared with the terminal branch lengths are 4.5 vs 60 for VNIa-4 (P-value from Kolmogorov-Smirnov test = 7×10^-70^), 3 vs 77.5 for VNIa-5 (P-value = 1×10^-53^) and 6 vs 44.5 for VNIa-93 (P-value = 4×10^-19^) (Supplementary Figure 6).

**Figure 2:**
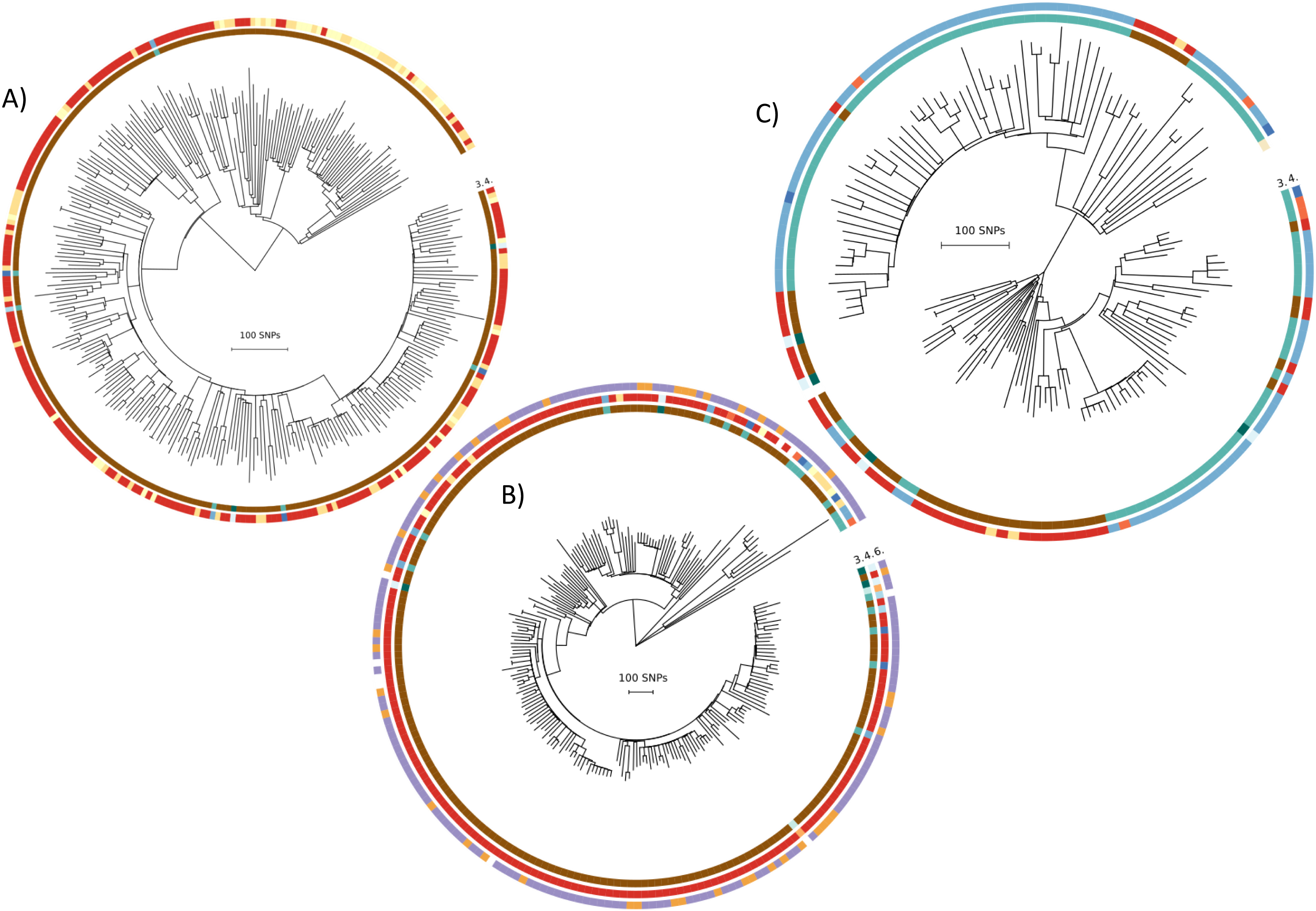
Within sub-clade phylogenetic trees for A) VNIa-4 B) VNIa-5 and C) VNIa-93. Rings are numbered and coloured according to Figure 1.

There were a total of 18071, 17593 and 7163 terminal branch SNPs in VNIa-4, VNIa-5 and VNIa-93. HIV infection status had no significant association with the terminal branch length of ST5 isolates. We had only 5 environmental strains in our dataset (one VNIa-4 and four VNIa-5), and they had a similar mean terminal branch length (75 SNPs). There were 263, 294 and 31 variants (1.5%, 1.8% and 0.4% of total) which occurred more than once on different terminal branches in VNIa-4, VNIa-5 and VNIa-93. However, most of these (VNIa-4, 52%; VNIa-5, 60%; and VNIa-93, 65%) were in intergenic regions (i.e. not in coding sequence, 3’ or 5’ UTR or introns). We manually investigated any gene containing a variant which occurred as a homoplasy in 3 or more strains for recognised links with virulence or host interactions, but had no informative hits. The average dN/dS of SNPs in the terminal branches were 0.84, 0.82 and 0.84 in VNIa-4, VNIa-5 and VNIa-93, respectively.

Another striking feature of the within sub-clade trees was the number of polytomies. All internal branches that represented 0 SNPs were collapsed, resulting in 78, 65 and 35 collapsed branches in 46, 36 and 21 distinct polytomies (defined as nodes with more than 2 children, after branches of 0 SNPs were collapsed) in VNIa-4, VNIa-5 and VNIa-93, respectively. The collapsed branches as a proportion of the total number of branches in each sub-clade were 13%, 15% and 12% in VNIa-4, VNIa-5 and VNIa-93. The median number of branches resulting from a polytomy event was 3 in all sub-clades, while the maximum was 9, 11 and 6 in VNIa-4, VNIa-5 and VNIa-93, respectively (Supplementary Table 4). For VNIa-4, 14 of 29 (48%) polytomies were international (i.e. strains in the polytomy were isolated from more than one country) and 1 (3%) of these was intercontinental. For VNIa-5, 10 of 24 (42%) polytomies were international and 6 (25%) were intercontinental. For VNIa-93, 4 of 21 (19%) polytomies were international and 1 (5%) of these was intercontinental. The maximum time separating the sampling date of two isolates descending directly from the same polytomy (i.e. not separated via an internal branch representing >0 SNPs) was 10 years for VNIa-4, 15 years for VNIa-5 and 8 years for VNIa-93. The median time range spanned by polytomies was 5.5, 5 and 1 year(s) for VNIa-4, VNIa-5 and VNIa-93, respectively. Genome sequences from isolates from both our study and that of Desjardins *et al.* belonged to the same polytomies.

### Within Sub-Clade Temporal Patterns

The majority of isolates in our study were collected during two clinical trials which recruited patients between 2004-2010 and 2013-2015 (Supplementary Figure 7A). As the first clinical trial only recruited patients in Vietnam, this is the only country for which we have considerable temporal range. This data shows that two sub-clades, VNIa-4 and VNIa-5 have been predominant in every year in which more than 5 samples were taken since 2004 (Supplementary Figure 7B). The prevalence of VNIa-32 appears to have declined, in 2004 it accounted for 12% (4/34) of *C. neoformans* collected, while there were no cases of this sub-clade observed in 2014 (0/40), the last year of collection. We found a lack of clock like evolution within all three sub-clades. The slope of the trend-line between time of isolation and root to tip distance was negative for both VNIa-4 and VNIa-5. There was a poor correlation between time of isolation and distance from the root in the tree for all three sub-clades (correlation co-efficient -0.07, -0.22 and 0.32 for VNIa-4, VNIa-5 and VNIa-93) (Supplementary Figure 8).

### Evidence of genome re-arrangement

The median number of genome re-arrangements between pairs of VNIa-4, VNIa-5 and VNIa-93 isolates were 10, 7 and 3, respectively. There was no significant association between SNP distance between isolates and the number of re-arrangements in VNIa-4, VNIa-5 or VNIa-93 (Supplementary Figure 9). There was also no association between the number of polytomies which occurred since the most recent common ancestor (MRCA) of the two isolates and the number of genome re-arrangements between the isolates (Supplementary Figure 10)

### Genome sequence and clinical features

#### Association between sub-clade and outcome

We used data from our recent randomised controlled trials of treatment for HIV-associated cryptococcal meningitis patients to define the effect of sub-clade on survival until 10 weeks or 6 months after randomisation. We used a Cox proportional hazards regression model with sub-clade as the main covariate, adjusted for country and treatment. Complete data were available from 530 patients. The survival over 6 months is illustrated in Figure 4. Infections with VNIa-93 were associated with a significantly reduced risk of death by both 10 weeks and 6 months (hazard ratios (HR) 0.45 95%CI 0.26 to 0.76, p = 0.003 and 0.60, 95%CI 0.39 to 0.94, p=0.024, respectively) compared with lineage VNIa-4 infections. There were no differences in outcomes between infections with VNIa-4 and any other lineage (See Supplementary Tables 5 and 6).

#### Association between VNIa-5 and HIV uninfected patients

Vietnam was the only country with more than 10 isolates of *C. neoformans* from HIV uninfected people. Therefore, only isolates from Vietnam were included in this analysis. Thirty five percent of HIV infected patients were infected with VNIa-5, compared with 75% of HIV uninfected patients (Fishers exact test, odds ratio 5.4, 95% CI 2.8-10.8, P < 10-8). Isolates from HIV uninfected patients are interspersed throughout the entire VNIa-5 phylogeny, implying all strains of this cluster could potentially cause infection in such hosts.

**Figure 4:**
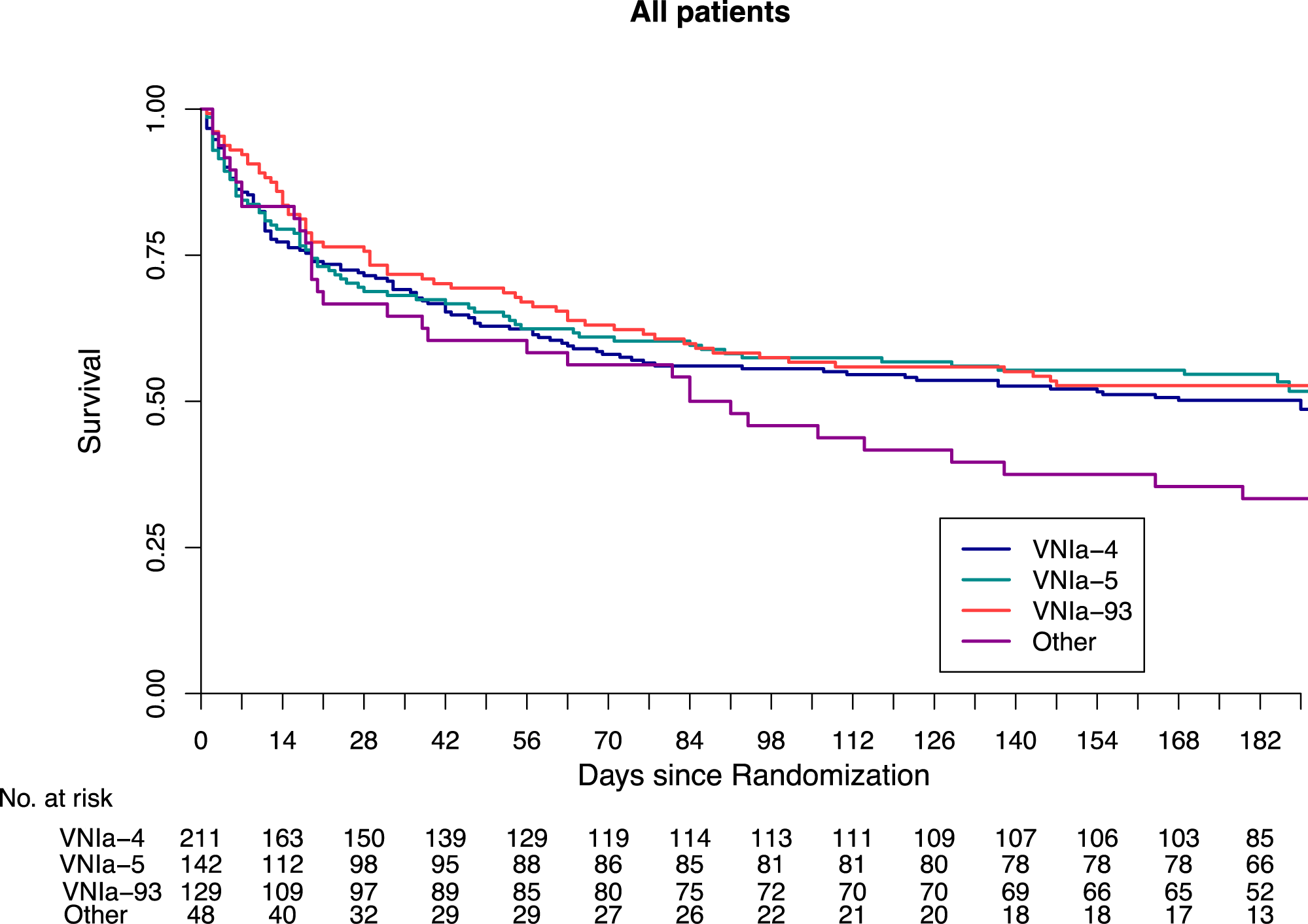
Kaplan-Meier survival estimates up to 6 months for all 530 HIV infected patients enrolled in one of two clinical trials (Day et al., 2013; Beardsley et al., 2016) with whole genome sequencing results for their infecting isolate.

#### VNIa-5 defining SNPs

Due to the association between VNIa-5 and disease in HIV uninfected patients, we were interested in SNPs which define VNIa-5. Ancestral sequence reconstruction identified 7465 SNPs between the ‘origin’ of VNIa-5 and the MRCA of VNIa-5 which were 95% sensitive and specific for VNIa-5. There were 1868 non-synonymous SNPs, distributed among 1220 genes. The dN/dS ratio was calculated for all genes with SNPs on the VNIa-5 defining branch, there were no genes known to be associated with virulence or interaction with the host that had extremes of dN/dS ratio. The overall dN/dS ratio of genic SNPs on this branch was 0.33, compared with the SNPs on the VNIa-4 defining branch which had an overall dN/dS of 0.38. There were seven genes with nonsense SNPs, introducing premature stop codons into five hypothetical proteins, one E3 ubiquitin-protein ligase (CNAG_04262) and a metacaspase, a cysteine protease involved in cell apoptosis (CNAG_06787).

### Mitochondrial sequence

A maximum likelihood phylogeny was derived for the SNPs identified in the mitochondrial DNA (mtSNP) of *C. neoformans* VNI (Supplementary Figure 11 B). When the mtSNP tree was compared with the whole genome SNP (wgSNP) tree (Supplementary Figure 11 B), some sub-clades were phylogenetically congruous, while others were not. VNIa-4, VNIa-5, VNIa-32, and VNIa-Y were all monophyletic within the mtSNP tree, in agreement with the whole genome SNP tree (Supplementary Figure 11 A). For VNIa-93, 144 out of 145 isolates were paraphyletic, with the monophyletic VNIa-32 and VNIa-Y nested within the VNIa-93 genotype, while VNIa-X was identical to the majority mtSNP genotype of VNIa-93. In the mitochondrial phylogeny VNIb is paraphyletic, giving rise to two sub-clades of VNIc, the first contained 19 isolates while the second is a singleton, and two VNI-outlier isolates. The most parsimonious description for VNIc is polyphyletic, with 8 different mono or paraphyletic groups. Otherwise, the paraphyletic grouping of all VNIc includes 648 isolates, only 89 of which are VNIc.

The most striking incongruity between the mtSNP and the whole genome data was in the placement of VNIa-5. In the whole genome tree, VNIa-5 is within the VNIa group with VNIa-4 as its sister taxa. In contrast, in the mtSNP tree, VNIa-5 is an outgroup, even in relation to VNIb and VNIc. There was a 28 bp sequence, intergenic between CNAG_09008 and CNAG_09009 (positions 19441 to 19469 of the mtSNP sequence, NC_018792.1), which contained 8 variants, present in every VNIa-5 in the dataset. This sequence begins 280 bp downstream of the 3’ end of CNAG_09008 and terminates 200 bp upstream of CNAG_09009. It had a per-site substitution rate of 0.28 compared with 0.004 for the VNIa-5 mitochondrial sequence as a whole. None of the variant positions were shared by any other *C. neoformans* strain, or by *C. deneoformans* JEC21 (GCA_000091045) or *C. gattii* R265 (GCA_000149475). When the putative recombinant region was compared against the full nr/nt BLAST database, the closest hit was to *C. neoformans* H99, chromosome 5 (NC_026749.1), positions 80207 to 80234, which had 1 bp difference (E-value = 0.004). This closest sequence on chromosome 5 is within CNAG_06848 which is widely conserved in the fungal kingdom. CNAG_06848 is a 222 bp gene encoding an ‘ATP synthase subunit 9, mitochondrial’. There were no strains in our dataset with SNPs in CNAG_06848 which could indicate a reciprocal recombination event. The assembly of the pacbio sequenced VNIa-5 genome also showed the presence of the highly variable region in the mitochondrial genome

## Discussion

We sequenced 699 isolates of *C. neoformans* covering 19 years and 5 countries on 2 continents, with most isolates derived from two large clinical trials. We integrated our novel data with previously published data (Desjardins et al. 2017) to provide extra context for our original findings. This context allowed us to assign 99.4% of the *C. neoformans* isolates sequenced as part of this study to the global clade VNI (Litvintseva et al. 2006; Khayhan et al. 2013; Ferreira-Paim et al. 2017). According to the nomenclature established by Desjardins *et al.* 98.5% of our isolates belonged to VNIa, compared with 30% of clinical VNI isolates and 18.5% of all isolates sequenced by Desjardins. To some extent, this difference is to be expected due to the focus of Desjardins *et al.* on both VNI and VNB, and their intensive sampling of Botswana, a known outlier in terms of *Cryptococcus* diversity (Litvintseva et al. 2006). This dominance of VNIa in our samples is interesting for two reasons. Firstly, it begs the question, are there specific biological properties of VNIa, or of VNIa-4, VNIa-5 and VNIa-93 which underlie their success? Secondly, the *C. neoformans* reference strain, H99, belongs to VNIb, which accounts for fewer than 1.5% of the clinical isolates in our study. We suggest that it may be useful to the *Cryptococcus* research community to consider including more representative isolates (i.e. from VNIa) in detailed laboratory investigations.

### There is very little novel diversity observed in the *C. neoformans* in our study

Even though 98.5% of our isolates were VNIa, we observed little additional diversity within VNIa that was not also observed in the much smaller number of VNIa isolates sequenced by Desjardins et al. This is due to the presence in our isolate collection of a small number of very common, highly clonal sub-clades. The three most common sub-clades (VNIa-4, VNIa-5 and VNIa-93) accounted for 92% of *C. neoformans* sequenced in this study. When there are a lot of internal nodes near the tips of the tree, it means that you either have high extinction rates or recently increased growth rate (Pybus et al. 2002). High extinction rate could be due to a relatively rapid decline in the ability of *C. neoformans* cells to germinate over time, while a recently increased growth rate could be due to exploitation of a new niche, such as the HIV infected human host.

### *C. neoformans* undergoes frequent transfers between continents

The phylo-geography of VNIa is characterised by each lineage being predominantly but not exclusively found in a single country or continent. While our sampling is exclusively from Asia and Africa, and is therefore not globally representative, VNIa-4 and VNIa-5 were predominantly Asian (97% and 89%), and VNIa-93 was predominantly African (64%). This finding is consistent with previous reports, with particular STs having been reported to be more common in certain countries, regions, or continents (Khayhan et al. 2013; Litvintseva et al. 2006; Ferreira-Paim et al. 2017). However, whole genome sequencing provides us with extra resolution in resolving whether, for example, the 7% of VNIa-5 strains in Africa are the result of a single introduction or multiple discrete introductions. To address this question, we generated within sub-clade reference genomes using PacBio sequencing and performed within sub-clade phylogenetic analyses. Examination of the within sub-clade phylogenetic trees (Figure 2) and parsimony analysis shows that international and intercontinental transmission is a frequent event, with 8-13% of internal branches representing an international transmission.

While nearly clonal isolates have been identified in disparate locations by a recent study (Rhodes et al. 2017), the authors focussed more on exploring ancient migrations. Our data dramatically illustrate the extent of this on-going intercontinental migration and we offer two alternative explanations. The first potential explanation is that transmission between countries or continents occurs during latent infection, i.e. a patient is exposed in one country, and then travels to another country where they develop illness and are sampled. Such long distance latent transmission has been hypothesised previously (Garcia-Hermoso et al. 1999). Unfortunately, we do not have extensive travel/residence histories for our patients and thus cannot directly address this hypothesis. However, historically there has not been large scale migration between Southeast Asia and South/East Africa (Kuyper 2008), suggesting that this hypothesis is insufficient to explain the high frequency of transmissions. A second, broad hypothesis to explain the large number of transmission events is that they are mediated by environmental factors, either ‘natural’ or human influenced. Potential natural environmental factors would include air currents or migratory birds; pigeons specifically are considered the most probable vector for global dissemination (Lin and Heitman 2006). Human activities that link the environments of East/Southeast Africa and Southeast Asia include trade in lumber, rice, exotic animals, and illegal animal products such as those used in traditional medicine e.g. ivory (http://www.aljazeera.com/news/2016/11/exclusive-vietnam-double-standards-ivory-trade-161114152646053.html). While we cannot directly address this hypothesis with our data, airborne spread is well established as a long distance dispersal mechanism for plant pathogens (Brown 2002). Intuitively it might seem unlikely that long distance airborne dispersal of fungal pathogens occurs frequently. However if airborne spore dispersal conforms to a non-exponentially bound (or ‘fat-tailed’) distribution model rather than an exponential model, long distance dispersions will occur relatively frequently (Brown 2002; Shaw 1994). Weather patterns are a proto-typical example of such ‘fat-tailed’, ‘chaotic’ (small differences in initial conditions, leading to large differences in outcome) distributions (Lorenz 1963). However, effective quantification of the potential contribution of airborne dispersal is complex (Meyer et al. 2017) and beyond the scope of this paper. Overall, we consider environmental factors to be the better explanation because (i) *Cryptococcus* is fundamentally an environmental organism (ii) there is limited human migration between Southeast Asia and East/Southeast Africa and (iii) long distance dispersal by environmental factors, including wind, is well established for fungal pathogens.

We found no correlation between root-to-tip phylogenetic distance and time since isolation in any of the three main VNIa sub-clades. One reason for the lack of a molecular clock could be the fact that *C. neoformans* can enter a quiescent state, in the form of hardy spores. The lack of molecular clock indicates *C. neoformans* spends enough time in the quiescent state to efface the clock-like signal, at least over the relatively short time scale sampled here. The lack of molecular clock has also been reported for the spore-forming bacterium *Bacillus anthracis* (Sahl et al. 2016).

### *C. neoformans* internal branches are very short compared with the terminal branches

A striking feature of all three within sub-clade phylogenies (VNIa-4, VNIa-5 and VNIa-93) was the difference between the length of the internal branches and the terminal branches. Median internal branch lengths were between 3 and 6 SNPs, while median terminal branch lengths were 44.5-77.5 SNPs. Long terminal branches are to be expected in an environmentally acquired organism, with no human-to-human transmission, but the contrast between these long terminal branches and the short internal branches is striking. Since the human host is considered a dead end for *C. neoformans* life-cycle, it is likely that most of the accumulation of variation represented by the internal branches occurred in the environment, rather than within humans. The short average internal branch indicates that *C. neoformans* VNIa does not generally acquire a lot of substitutions in the environment, implying a lack of cellular division and growth. In contrast, the terminal branches are almost all long, due to observed substitutions, and substitutions imply cell division. This leads to an apparent contradiction, as the terminal branches of the 5 environmental isolates in VNIa-4, VNIa-5 and VNIa-93 from our study and Desjardins et al. represent, on average, 75 SNPs. Therefore, to resolve this contradiction, we propose a scenario where *C. neoformans* VNIa is typically quiescent in the environment, and as a pre-condition for being cultured from the environment it must be recently derived from a population which is actively growing. This hypothesis could be tested by comparing the culture positive rates for *C. neoformans* from the environment, with positive rates by molecular testing e.g. PCR from the same samples. A higher number of positives for molecular testing compared with culture would support our hypothesis.

What are the implications of this idea for the clinical isolates which make up most of our cases? What we know is that the vast majority of clinical isolates have long terminal branches and that this amount of growth does not frequently occur in the environment. Therefore, the long terminal branches either occur within the patient, or are a pre-condition for infection of the human lung. We investigated the idea that, if the SNPs occur in patient the organisms may be evolving under pressure from the host, as has been observed for *C. neoformans* (Chen et al. 2017). The dN/dS of the SNPs in the terminal branches was consistent between sub-clades, ranging between 0.82-0.84 compared with the even lower dN/dS of SNPs in the long branches defining VNIa-4 (dN/dS = 0.38) and VNIa-5 (dN/dS = 0.33). This shows that SNPs in the terminal branches are relatively permissive of non-synonymous SNPs. This could be due to the SNPs occurring in an environment with relatively high positive selection, or more likely because there has been less time for mildly deleterious non-synonymous mutations to be lost. A small proportion (1.5%, 1.8%, 0.4%) of terminal branch SNPs were homoplasies. It is interesting that the majority of homoplasies were in intergenic regions, considering that according to standard models, these should be under weak or no selective pressure. It is possible that these intergenic regions have an unidentified regulatory role, as has recently been proposed for bacteria (Thorpe et al. 2017; Hammarlöf et al. 2018), or the homoplasies could be the result of recombination.

While we cannot put an accurate molecular clock to this dataset, it has been reported that within patients there is accumulation of 1 SNP every 58 days (Chen et al. 2017). If this rate holds for the terminal branches in our analysis then the average terminal branch represents between 7 and 12 years. This is an unfeasible length of time for a patient to have an uncontrolled *C. neoformans* infection, so either there is growth during latency or the substitution rate accounting for the terminal branch SNPs is higher than that reported by Chen *et al*. or much of the terminal branch mutation occurs outside the infected human.

### Large numbers of polytomies in *C. neoformans* phylogeny

One reason behind the short average internal branch length was the high number of polytomies in each sub-clade. A polytomy is a section of a phylogeny that cannot be fully resolved into dichotomous branching events. When observed in a phylogeny, they can either be due to a lack of information which allows the true relationship to be revealed (a ‘soft’ polytomy) or due to more than 2 simultaneous ‘speciation’ events (a ‘hard’ polytomy). As we are using reference genomes that are on average 277-361 SNPs from the isolates, in a ~19 Mbp genome, it seems unlikely that we are lacking SNP information that would resolve these polytomies. The lowest percentage coverage of the H99 reference genome in our analysis was 95%, indicating that the vast majority of the genome is being interrogated in these analyses. Therefore, the large numbers of polytomies we have observed are likely to be hard polytomies. Of course, at this fine scale, they are not speciation events, but rather the seeding of numerous progeny by a genetically homogenous population with no or very limited intermediate growth (if there was lots of intermediate growth, there would be accumulation of substitutions).

These polytomies occurred throughout the sub-clade trees, both near the tips and deeper in the tree. We showed that the isolates arising directly (i.e. immediate inferred ancestor was a polytomy) from the same polytomy could be remarkably diverse in their spatio-temporal distribution. The maximum difference in isolation time between two isolates arising from the same polytomy was 10 years, and the average ranged between 1 and 5.5 years for the different sub-clades. This temporal spread is not that surprising, considering the extended latent period of infection, and the lack of molecular clock in *C. neoformans*. What is more surprising is that 14-49% of polytomies result in infections of patients from different countries, and 3-25% result in infections of patients on different continents. This means that polytomies are unlikely to be entirely explained by exposure of all patients to the same point source of infectious propagules. One biological process which could explain these polytomies is long distance transmission of quiescent propagules. In other published studies, there were very few phylogenetically informative SNPs (two between 10 strains) reported in *Cryptococcus gattii* VGIIa, although the authors ascribed this to lack of sampling; it is unclear to us how the addition of further isolates will provide information which further differentiates these published sequences (Billmyre et al. 2014). Polytomies have been observed in phylogenetic analysis of *Bacillus anthracis* (Sahl et al. 2016), which also forms spores. A similar pattern of long terminal branches and short internal branches can be seen in the spore forming *C. difficile* (Knetsch et al. 2017).

Desjardins et al. established that there is still recombination on-going within VNIa. However, within each sub-clade, recombination appears to be a relatively minor contributor of genetic diversity. LD decay over genomic distance was minimal in all three sub-clades, although the small number of SNPs with a Minor Allele Frequency > 0.1 (due to short internal branches) means that this analysis was not well powered.

### Is the *C. neoformans* spore the quiescent propagule?

We present multiple strands of evidence (polytomies, difference between internal and terminal branch lengths, lack of molecular clock, long distance dispersal) which indicate that a quiescent phase is important in the epidemiology of *Cryptococcus neoformans*. The most obvious candidate for this quiescent stage is the well described *Cryptococcus neoformans* spore, which can be produced by either mating or fruiting. Due to the preponderance of one mating type, it is more likely that same sex fruiting is responsible for spore generation than mating (Lin et al. 2005). High rates of recombination have been reported during both fruiting and mating (Lin et al. 2005; Ni et al. 2013). However, in our data there was limited within sub-clade recombination, consistent with the lack of recombination within closely related outbreak clades of *C. gattii* in the Pacific Northwest (Billmyre et al. 2014). There was also limited genome re-arrangement, and little correlation between the number of SNPs and the number of genome re-arrangements.

Our failure to identify significant recombination leads us to believe that either fruiting is not the phenomenon which produces quiescent propagules or our results suffer from technical bias due to use of short read assemblies to detect genome re-arrangements. The *C. neoformans* genome is rich in transposons, which would be expected to break a short-read assembly, therefore transposon mediated re-arrangements (Idnurm et al. 2005) may not be detected by our analyses. Therefore, long read (Oxford Nanopore or Pacbio) sequencing should be used to address this technical explanation. Our data highlight an inconsistency in the literature between the clonal nature of globally distributed VNI and the putative role of spores in the natural history of cryptococcosis when recombination is expected to occur in both fruiting and mating. An alternative quiescent propagule which has been previously described in *C. neoformans* is desiccated yeast cells. Although minimal work has been done on this cell type, there is no obvious requirement for genetic re-arrangements during the desiccation process.

### Association between lineage and clinical features

We observed two associations between lineage and clinical phenotype. Firstly, the previously described association between VNIa-5 and the infection of HIV uninfected patients (Day et al. 2017) and secondly, the novel finding of a significantly lower risk of death at 10 weeks in patients infected with VNIa-93, in contrast to previous findings (Wiesner et al. 2012).

One interesting difference between the VNIa-5 isolates and the rest of VNIa was identified in the mitochondrial sequence. We observed a small recombination event which introduced 8 SNPs which were present in every VNIa-5 isolate and absent in every non-VNIa-5 isolate. The most likely candidate for the donor sequence was chromosome 5 of the *C. neoformans* nuclear genome, which encodes a sequence which varies by only 1 bp from the 21 bp putative recombinant fragment. While this has not been previously described in the literature, since these positions were not mixed, and the reads containing the divergent sequence mapped well to the mtDNA, we deem it likely that the mitochondrial sequence has been accurately re-constructed, while the origin of the divergent sequence is much less certain. That this change occurs in the mitochondrion is particularly intriguing as changes in mitochondrial morphology have been reported as underlying the hyper-virulence of the Vancouver outbreak *C. gattii* (Ma et al. 2009; Voelz et al. 2014). The putative recombination is in an intergenic region of the mitochondrion so if this variant underlies a modified phenotype, it is likely driven by changes in gene expression. Fungal mitochondrial 5’ untranslated leader sequences have been described between 81 to 220 bp in length (Schäfer 2005), while the putative recombination occurs 200 bp upstream of CNAG_09009.

In summary, the analysis of 699 *Cryptococcus* genomes has revealed that clinical isolates of *C. neoformans* from Vietnam, Laos, Thailand, Uganda and Malawi are concentrated in three main sub-clades. The phylogenetic structure indicates that there is either a high extinction rate in isolates causing human infections, or there has been a recent rapid expansion e.g. into a new niche such as HIV infected people. While it is frequently transmitted between continents, it likely spends the majority of it’s time in the environment in a quiescent state, but has always undergone a significant period of growth when cultured from the environment or infected people. We also show that VNIa-93, which has previously been associated with poorer outcomes is associated with a significantly reduced risk of death by 10 weeks compared with VNIa-4. We show that genome sequencing for fungal pathogens can provide insight into diverse clinical, epidemiological and ecological features.

## Materials and Methods (1188)

### Strain Selection

The Vietnamese isolates (N=441) were clinical isolates from the cerebrospinal fluid (CSF) of patients enrolled in a prospective, descriptive study of HIV-uninfected patients with central nervous system (CNS) infections (n=67) enrolled between 1997 and 2014, a randomized controlled trial of antifungal therapy in HIV-infected patients between 2004 and 2011, the CryptoDex trial, and 3 environmental isolates from Ho Chi Minh City, Vietnam (Beardsley et al. 2016; Chau et al. 2010; Day et al. 2011, 2013). The whole genome sequences of 8 Vietnamese strains in this analysis have been previously reported (Day et al. 2017). Lao isolates were from 73 patients with invasive cryptococcal infection admitted to Mahosot Hospital, Vientiane, between 2003 and 2015, including 5 from the CryptoDex trial. Isolates from Uganda (132), Malawi (13) and Thailand (40) were all from HIV infected patients enrolled into the CryptoDex trial (Beardsley et al. 2016). Sixty-nine isolates from Vietnam and 8 from Laos were derived from patients not known to be infected with HIV. All clinical trials had ethical approval from the local IRB in each centre and from the Oxford Tropical Ethics Committee.

### Micro and molecular biology

Isolates were revived from storage by incubation on Sabouraud’s agar at 30°C for 72 h. Single colonies were spread for confluent growth and incubated at 30°C for 24 h. For Illumina sequencing, genomic DNA was extracted from approximately 0.5 g (wet weight) of yeast cells using the MasterPure Yeast DNA purification kit (Epicentre, USA) according to manufacturer’s instructions. Whole genome sequencing was carried out on Illumina HiSeq 2000 at the Sanger Institute UK, and commercially through Macrogen, Korea using the HiSeq 4000 platform. DNA for PacBio sequencing was extracted according to the protocol in Supplementary methods which was modified from dx.doi.org/10.17504/protocols.io.ewtbfen. PacBio sequencing was performed by Macrogen, Seoul, Korea, for 20kb SMRT library production, with 2 SMRT cells per sample, according to the manufacturer’s instructions.

### Species identification, principal components analysis

Species identification was carried out using mash screen function (Ondov et al. 2016) comparing the sample FASTQs against the whole refseq database. For the principal components analysis all variant positions were loaded into an adegenet (Jombart and Ahmed 2011) (devel branch, commit 43b4360) genlight object using RStudio. Then the ade4 dudi.pca function was used to determine the principal components. K-means clustering was run on the first two principal components, with values to K between 2 and 10. The total within-cluster sum of squares was plotted for each K, and the number of clusters determined as the ‘elbow’ in the plot of K vs total within-cluster sum of squares. As the previously described VNIb and VNIc were grouped into one cluster in the analysis of the first two PCs, the same analysis was carried out on the 3rd and 4th PCs, which separated these two established lineages.

### Phylogenetics analysis

FASTQ data were mapped against the H99 reference (GCF_000149245) using bwa mem (Li 2013), SNPs were called using GATK v3.3.0 (McKenna et al. 2010) in unified genotyper mode. Positions where the majority allele accounted for < 90% of reads mapped at that position, which had a genotype quality of < 30, coverage < 5x, or mapping quality < 30 were recorded as Ns in further analyses. These steps were carried out using the PHEnix pipeline (https://github.com/phe-bioinformatics/PHEnix) and SnapperDB (Dallman et al. 2018). Positions in which at least one strain had a SNP passing quality thresholds were extracted and used as the input for RAxML v8.2.8 (Stamatakis 2014) maximum likelihood phylogenetic analysis. Ancestral state reconstruction was carried out using IQ-TREE v1.6 (Nguyen et al. 2015). To place our data into the broadest possible context, we included WGS data from Desjardins *et al*., 2017. To ensure efficient use of computational resources, a preliminary phylogenetic analysis was carried out, including all our data and representatives of VNI, VNBI, VNBII and VNII from Desjardins et al. For polytomy analysis ete3 (Huerta-Cepas et al. 2010) was used to delete/collapse nodes (branches) in the tree that represented 0 SNPs. Any node in this new tree with collapsed branches with 3 or more children was defined as a polytomy. Pacbio data was assembled using Canu v1.5 (Koren et al. 2017) and default parameters, polishing with Illumina data from the corresponding isolate using Pilon v1.22 (Walker et al. 2014) for multiple rounds until the number of indels being corrected per round was less than 2.

### Analysis of effect of sub-clade on outcome

We assessed the effect of sub-clade on time to death (10 weeks and 6 months) in HIV infected patients with cryptococcal meningitis with a Cox proportional hazards regression model with sub-clade as the main covariate. We included all patients with available data from our two randomized controlled trials. The model was adjusted for country, induction antifungal treatment (amphotericin monotherapy for 4 weeks, amphotericin combined with flucytosine for 2 weeks, or amphotericin combined with fluconazole for 2 weeks) and the use of adjunctive treatment with dexamethasone (Beardsley et al. 2016; Day et al. 2013). We tested the proportional hazard assumption based on scaled Schoenfeld residuals. Since we knew from the Cryptodex trial that the covariate dexamethasone does not satisfy this assumption, we included a time varying coefficient for dexamethasone use.

### Recombination analysis

Recombination analysis was carried out independently for VNIa-4, VNIa-5 and VNIa-93. Linkage disequilibrium (R2) was calculated on a per-lineage basis using vcftools v0.1.14 (Danecek et al. 2011) and the –geno-r2 option and a minimum allele frequency (MAF) of 0.1, LD was grouped in 100000 bp windows as there were not many SNPs with a MAF > 0.1 within sub-clades due to the short internal branches.

### Genome rearrangement analysis

Twenty five representatives of each sub-clade were *de novo* assembled using Velvet according to previously published methods (Makendi et al. 2016) and pairwise alignment carried out with Mauve (snapshot_2015-02-25) (Darling et al. 2010). The XMFA output of Mauve was then parsed using this python script (https://gist.github.com/flashton2003/b6c3e4e31e9084220fd30188988808f5) which briefly, looked within contigs with more than one co-linear block and checks whether the paired contig has more than one co-linear block. If so, it checks that the other co-linear blocks match between the two contigs and if not, infers a re-arrangement.

## Acknowledgements

JND was supported by a Wellcome Trust Intermediate fellowship WT097147MA. We would like to acknowledge the contribution of the Pathogen Informatics team at the Wellcome Sanger Institute, the Sequencing team at the Wellcome Sanger Institute, Macrogen of South Korea and MRC CLIMB for providing computational capacity (Connor et al. 2016). Isolates from Laos were obtained as part of the work programme of the Lao-Oxford-Mahosot Hospital Wellcome Trust Research Unit funded by the Wellcome Trust (106698/Z/14/Z). The authors are grateful to all the laboratory and clinical staff who helped with the collection of the isolates and data.

## References

Andrade-Silva LE, Ferreira-Paim K, Ferreira TB, Vilas-Boas A, Mora DJ, Manzato VM, Fonseca FM, Buosi K, Andrade-Silva J, Prudente da B S, et al. 2018. Genotypic analysis of clinical and environmental *Cryptococcus neoformans* isolates from Brazil reveals the presence of VNB isolates and a correlation with biological factors ed. K. Nielsen. PLoS One 13: e0193237. http://dx.plos.org/10.1371/journal.pone.0193237.

Beale MA, Sabiiti W, Robertson EJ, Fuentes-Cabrejo KM, O’Hanlon SJ, Jarvis JN, Loyse A, Meintjes G, Harrison TS, May RC, et al. 2015. Genotypic diversity is associated with clinical outcome and phenotype in cryptococcal meningitis across Southern Africa. PLoS Negl Trop Dis 9: 1–18.

Beardsley J, Wolbers M, Kibengo FM, Ggayi A-BM, Kamali A, Cuc NTK, Binh TQ, Chau NVV, Farrar J, Merson L, et al. 2016. Adjunctive Dexamethasone in HIV-Associated Cryptococcal Meningitis. N Engl J Med 374: 542–554. http://www.nejm.org/doi/10.1056/NEJMoa1509024.

Billmyre RB, Croll D, Li W, Mieczkowski P, Carter DA, Cuomo CA, Kronstad JW, Heitman J. 2014. Highly Recombinant VGII *Cryptococcus gattii* Population Develops Clonal Outbreak Clusters through both Sexual Macroevolution and Asexual Microevolution. MBio 5: e01494-14-e01494-14. http://mbio.asm.org/cgi/doi/10.1128/mBio.01494-14.

Brown JKM. 2002. Aerial Dispersal of Pathogens on the Global and Continental Scales and Its Impact on Plant Disease. Science (80-) 297: 537–541. http://www.sciencemag.org/cgi/doi/10.1126/science.1072678.

Chau TT, Mai NH, Phu NH, Nghia HD, Chuong L V, Sinh DX, Duong VA, Diep PT, Campbell JI, Baker S, et al. 2010. A prospective descriptive study of cryptococcal meningitis in HIV uninfected patients in Vietnam - high prevalence of *Cryptococcus neoformans* var *grubii* in the absence of underlying disease. BMC Infect Dis 10: 199. http://bmcinfectdis.biomedcentral.com/articles/10.1186/1471-2334-10-199.

Chen Y, Farrer RA, Giamberardino C, Sakthikumar S, Jones A, Yang T, Tenor JL, Wagih O, Van Wyk M, Govender NP, et al. 2017. Microevolution of Serial Clinical Isolates of *Cryptococcus neoformans* var. *grubii* and *C. gattii* ed. F. Dromer. MBio 8: e00166-17. http://mbio.asm.org/lookup/doi/10.1128/mBio.00166-17.

Connor TR, Loman NJ, Thompson S, Smith A, Southgate J, Poplawski R, Bull MJ, Richardson E, Ismail M, Thompson SE-, et al. 2016. CLIMB (the Cloud Infrastructure for Microbial Bioinformatics): an online resource for the medical microbiology community. Microb Genomics 2. http://www.microbiologyresearch.org/content/journal/mgen/10.1099/mgen.0.000086.

Dallman T, Ashton P, Schafer U, Jironkin A, Painset A, Shaaban S, Hartman H, Myers R, Underwood A, Jenkins C, et al. 2018. SnapperDB: a database solution for routine sequencing analysis of bacterial isolates. Bioinformatics 81: 3946–3952. https://academic.oup.com/bioinformatics/advance-article/doi/10.1093/bioinformatics/bty212/4961427.

Danecek P, Auton A, Abecasis G, Albers CA, Banks E, DePristo MA, Handsaker RE, Lunter G, Marth GT, Sherry ST, et al. 2011. The variant call format and VCFtools. Bioinformatics 27: 2156–2158. https://academic.oup.com/bioinformatics/article-lookup/doi/10.1093/bioinformatics/btr330.

Darling AE, Mau B, Perna NT. 2010. progressiveMauve: Multiple Genome Alignment with Gene Gain, Loss and Rearrangement ed. J.E. Stajich. PLoS One 5: e11147. http://dx.plos.org/10.1371/journal.pone.0011147.

Day JN, Chau TTH, Wolbers M, Mai PP, Dung NT, Mai NH, Phu NH, Nghia HD, Phong ND, Thai CQ, et al. 2013. Combination Antifungal Therapy for Cryptococcal Meningitis. N Engl J Med 368: 1291–1302. http://www.nejm.org/doi/10.1056/NEJMoa1110404.

Day JN, Hoang TN, Duong A V, Hong CTT, Diep PT, Campbell JI, Sieu TPM, Hien TT, Bui T, Boni MF, et al. 2011. Most Cases of Cryptococcal Meningitis in HIV-Uninfected Patients in Vietnam Are Due to a Distinct Amplified Fragment Length Polymorphism-Defined Cluster of Cryptococcus neoformans var. grubii VN1. J Clin Microbiol 49: 658–664. http://jcm.asm.org/cgi/doi/10.1128/JCM.01985-10.

Day JN, Qihui S, Thanh LT, Trieu PH, Van AD, Thu NH, Chau TTH, Lan NPH, Chau NVV, Ashton PM, et al. 2017. Comparative genomics of *Cryptococcus neoformans* var. *grubii* associated with meningitis in HIV infected and uninfected patients in Vietnam ed. J.M. Vinetz. PLoS Negl Trop Dis 11: e0005628. http://dx.plos.org/10.1371/journal.pntd.0005628%5Cn http://www.ncbi.nlm.nih.gov/pubmed/28614360.

Desjardins CA, Giamberardino C, Sykes SM, Yu C-H, Tenor JL, Chen Y, Yang T, Jones AM, Sun S, Haverkamp MR, et al. 2017. Population genomics and the evolution of virulence in the fungal pathogen *Cryptococcus neoformans*. Genome Res 118323. http://genome.cshlp.org/lookup/doi/10.1101/gr.218727.116.

Ferreira-Paim K, Andrade-Silva L, Fonseca FM, Ferreira TB, Mora DJ, Andrade-Silva J, Khan A, Dao A, Reis EC, Almeida MTG, et al. 2017. MLST-Based Population Genetic Analysis in a Global Context Reveals Clonality amongst *Cryptococcus neoformans* var. *grubii* VNI Isolates from HIV Patients in Southeastern Brazil. PLoS Negl Trop Dis 11: e0005223. http://dx.plos.org/10.1371/journal.pntd.0005223.

Garcia-Hermoso D, Janbon G, Dromer F. 1999. Epidemiological evidence for dormant *Cryptococcus neoformans* infection. J Clin Microbiol 37: 3204–9. http://www.ncbi.nlm.nih.gov/pubmed/10488178.

Hammarlöf DL, Kröger C, Owen S V., Canals R, Lacharme-Lora L, Wenner N, Schager AE, Wells TJ, Henderson IR, Wigley P, et al. 2018. Role of a single noncoding nucleotide in the evolution of an epidemic African clade of Salmonella. Proc Natl Acad Sci 115: E2614–E2623. http://www.pnas.org/lookup/doi/10.1073/pnas.1714718115.

Hiremath SS, Chowdhary A, Kowshik T, Randhawa HS, Sun S, Xu J. 2008. Long-distance dispersal and recombination in environmental populations of *Cryptococcus neofarmans* var. *grubii* from India. Microbiology 154: 1513–1524.

Huerta-Cepas J, Dopazo J, Gabaldón T. 2010. ETE: a python Environment for Tree Exploration. BMC Bioinformatics 11: 24. http://bmcbioinformatics.biomedcentral.com/articles/10.1186/1471-2105-11-24.

Idnurm A, Bahn Y-S, Nielsen K, Lin X, Fraser JA, Heitman J. 2005. Deciphering the Model Pathogenic Fungus *Cryptococcus Neoformans*. Nat Rev Microbiol 3: 753–764. http://www.nature.com/doifinder/10.1038/nrmicro1245.

Jombart T, Ahmed I. 2011. adegenet 1.3-1: New tools for the analysis of genome-wide SNP data. Bioinformatics 27: 3070–3071.

Kaocharoen S, Ngamskulrungroj P, Firacative C, Trilles L, Piyabongkarn D, Banlunara W, Poonwan N, Chaiprasert A, Meyer W, Chindamporn A. 2013. Molecular Epidemiology Reveals Genetic Diversity amongst Isolates of the *Cryptococcus neoformans*/*C. gattii* Species Complex in Thailand ed. B. Wanke. PLoS Negl Trop Dis 7: e2297. http://dx.plos.org/10.1371/journal.pntd.0002297.

Khayhan K, Hagen F, Pan W, Simwami S, Fisher MC, Wahyuningsih R, Chakrabarti A, Chowdhary A, Ikeda R, Taj-Aldeen SJ, et al. 2013. Geographically Structured Populations of *Cryptococcus neoformans* Variety *grubii* in Asia Correlate with HIV Status and Show a Clonal Population Structure. PLoS One 8: 1–14.

Knetsch CW, Kumar N, Forster SC, Connor TR, Browne HP, Harmanus C, Sanders IM, Harris SR, Turner L, Morris T, et al. 2017. Zoonotic Transfer of *Clostridium difficile* Harboring Antimicrobial Resistance between Farm Animals and Humans ed. B. Fenwick. J Clin Microbiol 56: e01384-17. http://jcm.asm.org/lookup/doi/10.1128/JCM.01384-17.

Koren S, Walenz BP, Berlin K, Miller JR, Bergman NH, Phillippy AM. 2017. Canu: scalable and accurate long-read assembly via adaptive k -mer weighting and repeat separation. Genome Res 27: 722–736. http://genome.cshlp.org/lookup/doi/10.1101/gr.215087.116.

Kuyper M. 2008. Return Migration to Vietnam.

Li H. 2013. Aligning sequence reads, clone sequences and assembly contigs with BWA-MEM. arXiv. http://arxiv.org/abs/1303.3997.

Lin X, Heitman J. 2006. The biology of the *Cryptococcus neoformans* species complex. AnnuRevMicrobiol 60: 69–105.

Lin X, Hull CM, Heitman J. 2005. Sexual reproduction between partners of the same mating type in *Cryptococcus neoformans*. Nature 434: 1017–1021. http://www.nature.com/doifinder/10.1038/nature03448.

Litvintseva AP, Thakur R, Vilgalys R, Mitchell TG. 2006. Multilocus sequence typing reveals three genetic subpopulations of *Cryptococcus neoformans* var. *grubii* (serotype A), including a unique population in Botswana. Genetics 172: 2223–2238.

Lorenz EN. 1963. Deterministic Nonperiodic Flow. J Atmos Sci 20: 130–141. http://journals.ametsoc.org/doi/abs/10.1175/1520-0469%281963%29020%3C0130%3ADNF%3E2.0.CO%3B2.

Ma H, Hagen F, Stekel DJ, Johnston SA, Sionov E, Falk R, Polacheck I, Boekhout T, May RC. 2009. The fatal fungal outbreak on Vancouver Island is characterized by enhanced intracellular parasitism driven by mitochondrial regulation. Proc Natl Acad Sci 106: 12980–12985. http://www.pnas.org/cgi/doi/10.1073/pnas.0902963106.

Makendi C, Page AJ, Wren BW, Le Thi Phuong T, Clare S, Hale C, Goulding D, Klemm EJ, Pickard D, Okoro C, et al. 2016. A Phylogenetic and Phenotypic Analysis of Salmonella enterica Serovar Weltevreden, an Emerging Agent of Diarrheal Disease in Tropical Regions ed. E.T. Ryan. PLoS Negl Trop Dis 10: e0004446. http://dx.plos.org/10.1371/journal.pntd.0004446.

McKenna A, Hanna M, Banks E, Sivachenko A, Cibulskis K, Kernytsky A, Garimella K, Altshuler D, Gabriel S, Daly M, et al. 2010. The Genome Analysis Toolkit: A MapReduce framework for analyzing next-generation DNA sequencing data. Genome Res 20: 1297–1303. http://genome.cshlp.org/cgi/doi/10.1101/gr.107524.110.

Meyer M, Cox JA, Hitchings MDT, Burgin L, Hort MC, Hodson DP, Gilligan CA. 2017. Quantifying airborne dispersal routes of pathogens over continents to safeguard global wheat supply. Nat Plants 3: 780–786. http://www.nature.com/articles/s41477-017-0017-5.

Nguyen L, Schmidt HA, von Haeseler A, Minh BQ. 2015. IQ-TREE: A Fast and Effective Stochastic Algorithm for Estimating Maximum-Likelihood Phylogenies. Mol Biol Evol 32: 268–274. https://academic.oup.com/mbe/article-lookup/doi/10.1093/molbev/msu300.

Ni M, Feretzaki M, Li W, Floyd-Averette A, Mieczkowski P, Dietrich FS, Heitman J. 2013. Unisexual and Heterosexual Meiotic Reproduction Generate Aneuploidy and Phenotypic Diversity De Novo in the Yeast *Cryptococcus neoformans* ed. A.P. Mitchell. PLoS Biol 11: e1001653. http://dx.plos.org/10.1371/journal.pbio.1001653.

Ondov BD, Treangen TJ, Melsted P, Mallonee AB, Bergman NH, Koren S, Phillippy AM. 2016. Mash: fast genome and metagenome distance estimation using MinHash. Genome Biol 17: 132. http://dx.doi.org/10.1186/s13059-016-0997-x.

Park BJ, Wannemuehler KA, Marston BJ, Govender N, Pappas PG, Chiller TM. 2009. Estimation of the current global burden of cryptococcal meningitis among persons living with HIV/AIDS. >AIDS 23: 525–530. http://content.wkhealth.com/linkback/openurl?sid=WKPTLP:landingpage&an=00002030-200902200-00012.

Pybus OG, Rambaut A, Holmes EC, Harvey PH. 2002. New inferences from tree shape: numbers of missing taxa and population growth rates. Syst Biol 51: 881–8. http://www.ncbi.nlm.nih.gov/pubmed/12554454.

Rajasingham R, Smith RM, Park BJ, Jarvis JN, Govender NP, Chiller TM, Denning DW, Loyse A, Boulware DR. 2017. Global burden of disease of HIV-associated cryptococcal meningitis: an updated analysis. Lancet Infect Dis 17: 873–881. http://dx.doi.org/10.1016/S1473-3099(17)30243-8.

Rhodes J, Desjardins CA, Sykes SM, Beale MA, Vanhove M, Sakthikumar S, Chen Y, Gujja S, Saif S, Chowdhary A, et al. 2017. Tracing Genetic Exchange and Biogeography of *Cryptococcus neoformans* var. *grubii* at the Global Population Level. Genetics 207: 327–346. http://www.genetics.org/lookup/doi/10.1534/genetics.117.203836.

Sahl JW, Pearson T, Okinaka R, Schupp JM, Gillece JD, Heaton H, Birdsell D, Hepp C, Fofanov V, Noseda R, et al. 2016. A *Bacillus anthracis* Genome Sequence from the Sverdlovsk 1979 Autopsy Specimens. MBio 7: e01501-16. http://mbio.asm.org/lookup/doi/10.1128/mBio.01501-16.

Schäfer B. 2005. RNA maturation in mitochondria of *S. cerevisiae* and S. pombe. Gene 354: 80–85. http://linkinghub.elsevier.com/retrieve/pii/S037811190500168X.

Shaw MW. 1994. Modeling Stochastic Processes in Plant Pathology. Annu Rev Phytopathol 32: 523–544. http://www.annualreviews.org/doi/10.1146/annurev.py.32.090194.002515.

Simwami SP, Khayhan K, Henk DA, Aanensen DM, Boekhout T, Hagen F, Brouwer AE, Harrison TS, Donnelly CA, Fisher MC. 2011. Low Diversity *Cryptococcus neoformans* Variety *grubii* Multilocus Sequence Types from Thailand Are Consistent with an Ancestral African Origin ed. J. Heitman. PLoS Pathog 7: e1001343. http://dx.plos.org/10.1371/journal.ppat.1001343.

Stamatakis A. 2014. RAxML version 8: a tool for phylogenetic analysis and post-analysis of large phylogenies. Bioinformatics 30: 1312–1313. https://academic.oup.com/bioinformatics/article-lookup/doi/10.1093/bioinformatics/btu033.

Thanh LT, Trieu PH, Rattanavong S, Trinh MN, Anh D Van, Dacon C, Thu HN, Lan PHN, Chau THT, Davong V, et al. 2017. Multilocus Sequence Typing Reveals a Unique Co-dominant Population Structure of *Cryptococcus neoformans* var. *grubii* in Vietnam. bioRxiv.

Thorpe HA, Bayliss SC, Hurst LD, Feil EJ. 2017. Comparative Analyses of Selection Operating on Nontranslated Intergenic Regions of Diverse Bacterial Species. Genetics 206: 363–376. http://www.genetics.org/lookup/doi/10.1534/genetics.116.195784.

Vanhove M, Beale MA, Rhodes J, Chanda D, Lakhi S, Kwenda G, Molloy S, Karunaharan N, Stone N, Harrison TS, et al. 2017. Genomic epidemiology of Cryptococcus yeasts identifies adaptation to environmental niches underpinning infection across an African HIV/AIDS cohort. Mol Ecol 26: 1991–2005. http://doi.wiley.com/10.1111/mec.13891.

Velagapudi R, Hsueh Y-P, Geunes-Boyer S, Wright JR, Heitman J. 2009. Spores as Infectious Propagules of *Cryptococcus neoformans*. Infect Immun 77: 4345–4355. http://iai.asm.org/cgi/doi/10.1128/IAI.00542-09.

Voelz K, Johnston SA, Smith LM, Hall RA, Idnurm A, May RC. 2014. ‘Division of labour’ in response to host oxidative burst drives a fatal *Cryptococcus gattii* outbreak. Nat Commun 5: 5194. http://www.nature.com/doifinder/10.1038/ncomms6194.

Walker BJ, Abeel T, Shea T, Priest M, Abouelliel A, Sakthikumar S, Cuomo CA, Zeng Q, Wortman J, Young SK, et al. 2014. Pilon: An Integrated Tool for Comprehensive Microbial Variant Detection and Genome Assembly Improvement ed. J. Wang. PLoS One 9: e112963. http://dx.plos.org/10.1371/journal.pone.0112963.

Wiesner DL, Moskalenko O, Corcoran JM, McDonald T, Rolfes MA, Meya DB, Kajumbula H, Kambugu A, Bohjanen PR, Knight JF, et al. 2012. Cryptococcal Genotype Influences Immunologic Response and Human Clinical Outcome after Meningitis. MBio 3: e00196-12-e00196-12. http://mbio.asm.org/cgi/doi/10.1128/mBio.00196-12.

